# A global perspective on microbial diversity in the terrestrial deep subsurface

**DOI:** 10.1101/602672

**Authors:** A. Soares, A. Edwards, D. An, A. Bagnoud, M. Bomberg, K. Budwill, S. M. Caffrey, M. Fields, J. Gralnick, V. Kadnikov, L. Momper, M. Osburn, J.W. Moreau, D. Moser, A. Mu, L. Purkamo, S. M. Rassner, C. S. Sheik, B. Sherwood Lollar, B. M. Toner, G. Voordouw, K. Wouters, A. C. Mitchell

**Affiliations:** Department of Geography and Earth Sciences (DGES), Aberystwyth University (AU), Wales, UK; Institute of Biology, Environmental and Rural Sciences (IBERS), AU; Interdisciplinary Centre for Environmental Microbiology (iCEM), AU; Department of Biological Sciences, University of Calgary, Canada; Institut de Génie Thermique (IGT), Haute École d’Ingénierie et de Gestion du Canton de Vaud (HEIG-VD), Yverdon-les-Bains, Switzerland; VTT Technical Research Centre of Finland, Finland; Alberta Innovates, Canada; University of Toronto, Canada (UT); Center for Biofilm Engineering (CBE), Montana State University (MSU), USA; Department of Plant and Microbial Biology, UM, USA; Faculty of Biology, Moscow State University (MoSU), Russia; Institute of Bioengineering, Research Center of Biotechnology, Russian Academy of Sciences, Russia; Department of Earth, Atmospheric and Planetary Sciences (DEAPS), The Massachusetts Institute of Technology (MIT), United States of America (USA); Department of Earth and Planetary Sciences (DEPS), Northwestern University (NWU), USA; School of Earth Sciences, The University of Melbourne (UM), Parkville, Australia; Department of Microbiology and Immunology at the Peter Doherty Institute for Infection and Immunity, UM; Doherty Applied Microbial Genomics, Department of Microbiology and Immunology at the Peter Doherty Institute for Infection and Immunity, UM; Microbiological Diagnostic Unit Public Health Laboratory, Department of Microbiology and Immunology, UM; Division of Hydrologic Sciences, Desert Research Institute (DRI), Las Vegas, NV, USA; School of Earth and Environmental Sciences (SEES), University of St. Andrews (USA), Scotland, UK; Geological Survey of Finland (GTK), Finland; Large Lakes Observatory, University of Minnesota – Duluth (UMD); Department of Earth Sciences, UT, Canada; Department of Soil, Water & Climate, University of Minnesota; Institute for Environment, Health and Safety (EHS), Belgian Nuclear Research Centre SCK•CEN, Mol, Belgium

## Abstract

While recent efforts to catalogue Earth’s microbial diversity have focused upon surface and marine habitats, 12% to 20% of Earth’s bacterial and archaeal biomass is suggested to inhabit the terrestrial deep subsurface, compared to ∼1.8% in the deep subseafloor ^1–3^. Metagenomic studies of the terrestrial deep subsurface have yielded a trove of divergent and functionally important microbiomes from a range of localities ^4–6^. However, a wider perspective of microbial diversity and its relationship to environmental conditions within the terrestrial deep subsurface is still required. Here, we show the diversity of bacterial communities in deep subsurface groundwater is controlled by aquifer lithology globally, by using 16S rRNA gene datasets collected across five countries on two continents and from fifteen rock types over the past decade. Furthermore, our meta-analysis reveals that terrestrial deep subsurface microbiota are dominated by Betaproteobacteria, Gammaproteobacteria and Firmicutes, likely as a function of the diverse metabolic strategies of these taxa. Despite this similarity, evidence was found not only for aquifer-specific microbial communities, but also for a common small consortium of prevalent Betaproteobacteria and Gammaproteobacterial OTUs across the localities. This finding implies a core terrestrial deep subsurface community, irrespective of aquifer lithology, that may play an important role in colonising and sustaining microbial habitats in the deep terrestrial subsurface. An *in-silico* contamination-aware approach to analysing this dataset underscores the importance of downstream methods for assuring that robust conclusions can be reached from deep subsurface-derived sequencing data. Understanding the global panorama of microbial diversity and ecological dynamics in the deep terrestrial subsurface provides a first step towards understanding the role of microbes in global subsurface element and nutrient cycling.

## Main text

Understanding the distribution of microbial diversity is pivotal for advancing our knowledge of deep subsurface global biogeochemical cycles ^7, 8^. Subsurface biomass is suggested to have exceeded that of the Earth’s surface by an order of magnitude (∼45% of Earth’s total biomass) before land plants evolved, at ca. 0.5 billion years ago ^9^. Integrative modelling of cell count and quantitative PCR (qPCR) data and geophysical factors indicated in late 2018 that the bacterial and archaeal biomass found in the global deep subsurface may range from 23 to 31 petagrams of carbon (PgC). These values halved previous efforts from earlier that year^10^ but maintained the notion that the terrestrial deep subsurface holds ca. 5-fold more bacterial and archaeal biomass than the deep marine subsurface. Further, it is expected that 20-80% of the possible 2-6 × 10^29^ prokaryotic cells present in the terrestrial subterranean biome exist as biofilms and play crucial roles in global biogeochemical cycles ^10, 11^.

Cataloguing microbial diversity and functionality in the terrestrial deep subsurface has mostly been achieved by means of marker gene and metagenome sequencing in coals, sandstones, carbonates, and clays, as well as deep igneous and metamorphic rocks ^4–6, 12–20^. Only recently has the first comprehensive database of 16S rRNA gene-based studies targeting terrestrial subsurface environments been compiled ^10^. This work focused on updating estimates for bacterial and archaeal biomass, and cell numbers across the terrestrial deep subsurface, but also linked the identified bacterial and archaeal phylum-level compositions to host-rock type, and to 16S rRNA gene region primer targets ^10^. While highlighting Firmicutes and Proteobacterial dominance in the bacterial component of terrestrial deep subsurface, no further taxonomic insights were gained. However, genus-level identification is critical for understanding community composition, inferred metabolism and hence microbial contributions of distinct community members to biogeochemical cycling in the deep subsurface ^18, 21–23^. Indeed, such genus-specific traits have been demonstrated as critical for understanding crucial biological functions in other microbiomes ^24^, and genus-specific functions of relevance for deep subsurface biogeochemistry are clear ^25, 26^.

So far, the potential biogeochemical impacts of microbial activity in the deep subsurface have been inferred through shotgun metagenomics, as well as from incubation experiments of primary geological samples amended with molecules or minerals of interest ^13, 19, 20, 27–30^. Recent studies of deep terrestrial subsurface microbial communities further suggest that these are metabolically active, generally associated with novel uncultured phyla, and potentially directly involved in carbon and sulphur cycling ^31–36^. Concomitant advancements in subsurface drilling, molecular methods and computational techniques have aided the exploration of the subsurface biosphere, but serious challenges remain mostly related to deciphering sample contamination by drilling methods and sample transportation to laboratories for processing ^37, 38^. The logistical challenges inherent to accessing and recovering *in situ* samples from hundreds to thousands of metres below surface complicate our view of terrestrial subsurface microbial ecology ^39^.

In this study, we capitalize on the increased availability of 16S rRNA gene amplicon data from multiple studies of the terrestrial deep subsurface conducted over the last decade. We apply bespoke bioinformatic scripts to generate insights into the microbial community structure and controls upon bacterial microbiomes of the terrestrial deep subsurface across a large distribution of habitat types on multiple continents. The deep biosphere is as-yet undefined as a biome - elevated temperature, anoxic conditions, low levels of organic carbon, and measures of isolation from the surface photosphere are some of the criteria used albeit without a consensus. For this work a more general approach has been taken to define the terrestrial deep subsurface as the zone at least 100 m from the surface ^40, 41^.

### Meta-analysis of the terrestrial deep subsurface microbiome

Here, we were able to compare datasets encompassing different 16S rRNA gene hyper-variable regions, and derived from different DNA extraction methodologies, facilitated by closed-reference Operational Taxonomic Unit (OTU)-picking of each study individually using the same 16S rRNA gene reference database. This procedure begins to address technically confounding variables by limiting taxonomy assignments to only the archaeal and bacterial diversity listed in the chosen database and precludes the discovery of novel taxa.

The finalized meta-analysis dataset comprised of 16S rRNA data from seventeen aquifers in either sedimentary- or crystalline-host rocks, from depths spanning 94 m to 2300 m below land surface (mbls), targeting mostly groundwater across 5 countries and two continents (**Supplementary Table 3**). Nine DNA extraction techniques were used in these studies, ranging from standard and modified kit protocols (e.g. MOBIO^®^ PowerSoil, see **Table 1**) to phenol-chloroform and CTAB/NaCl based methods ^42–47^. Finally, 6 different primer pair amplified regions of the 16S rRNA gene, in 454 pyrosequencing and Illumina sequencing, were used to generate the datasets.

**Table 1.** Metadata table for the studies utilized in this meta-analysis (cf. Supplementary Table 3 and Supplementary Figure 3 for more details). NA is used as an acronym for “not available”. The dataset unavailable through SRA is available through http://hmp.ucalgary.ca/HMP/.

| SRA Accession | Originator | DOI | Year of Sampling | Final no. of samples | Location | Depth gradient (m) | Host-rock | Final no. of sequences | Final no. of OTUs |
| --- | --- | --- | --- | --- | --- | --- | --- | --- | --- |
| PRJNA262938 | Magdalena Osburn, Lily Momper | 10.3389/fmicb.2014.00610 | 2013/2014 | 6 | South Dakota, USA | 243.84-1478.28 | Sulphide-rich schists | 170364 | 1367 |
| PRJNA268940 | Duane Moser | NA | 2007-2011 | 8 | California, USA; Ontario, Canada | 94-2383 | Dolomite, Tuff, Rhyolite/tuff-breccia | 68071 | 741 |
| PRJNA248749 | Jeffrey Gralnick | NA | 2011 | 6 | Minnesota, USA | 730 | Hematite | 69757 | 511 |
| PRJNA251746 | Rick Colwell | NA | 2009 | 6 | Washington, USA | 393-1135 | Basalt | 30121 | 613 |
| PRJNA375701 | Elliot Barnhart | 10.1016/j.coal.2016.05.001 | 2013 | 5 | Montana, USA | 109-114.7 | Sub-bituminous coal, Shale, Sandstone, Siltstone | 35926 | 154 |
| NA | Karen Budwill | 10.1128/AEM.01737-15 | 2009 | 38 | Alberta, Canada | 140-1064.4 | Sub-bituminous coal, Volatile bituminous coal | 100618 | 5110 |
| PRJEB1468 | Katinka Wouters | 10.1111/1574-6941.12171 |  | 6 | Mol, Belgium | 217-232 | Kaolinite/illite and smectite | 47123 | 497 |
| PRJEB10822 | Lotta Purkamo | 10.5194/bg-13-3091-2016 | 2009-2011 | 7 | Outokumpu, Finland | 180-2300 | Mica-schist, Biotite-gneiss, Chlorite-sericite schist | 12290 | 177 |

Initial processing of 187 retrieved samples revealed 24,632,035 chimera-checked sequences ^17, 28, 43, 44, 48–50^. SILVA 123-aided closed-reference OTU-picking yielded 6,975 OTUs associated to 598,341 sequences following exclusion of singleton OTUs and samples containing 2 or less OTUs. The final dataset following stricter contamination-aware filtering (*cf.* **Methodology**) was comprised of 70 samples and 2,207 OTUs (513,929 sequences, 2.54% of the initial sequences), where Archaeal reads comprised 1.5% of the total number of reads.

### Trends in taxonomic diversity

Among a total of 45 detected bacterial phyla, Proteobacteria were seen to dominate most community profiles in this dataset (**Figure 1**). The most abundant proteobacterial classes (Alpha-, Betaproteobacteria, Delta-, Gammaproteobacteria) represented 57.2% of the total number of reads, with 13.4% of these assigned to class Clostridia (Firmicutes). A general prevalence of Betaproteobacteria and Gammaproteobacteria in the deep biosphere may be explained by the diverse metabolic capabilities of taxa within these clades. Families Gallionellaceae, Pseudomonadaceae, Rhodocyclaceae and Hydrogeniphillaceae within Betaproteobacteria and Gammaproteobacteria are suggested to play crucial roles in deep subsurface iron, nitrogen, sulphur and carbon cycling across the world ^43, 51, 52^. The relative abundance of order Burkholderiales (Betaproteobacteria) in surficial soils has previously been correlated (R^2^=0.92, ANOVA p-value <0.005) with mineral dissolution rates, while genus *Pseudomonas* (Gammaproteobacteria) is widely known to playing a key role in hydrocarbon-degradation, denitrification and coal solubilisation in different locations^53–55^.

**Figure 1.**
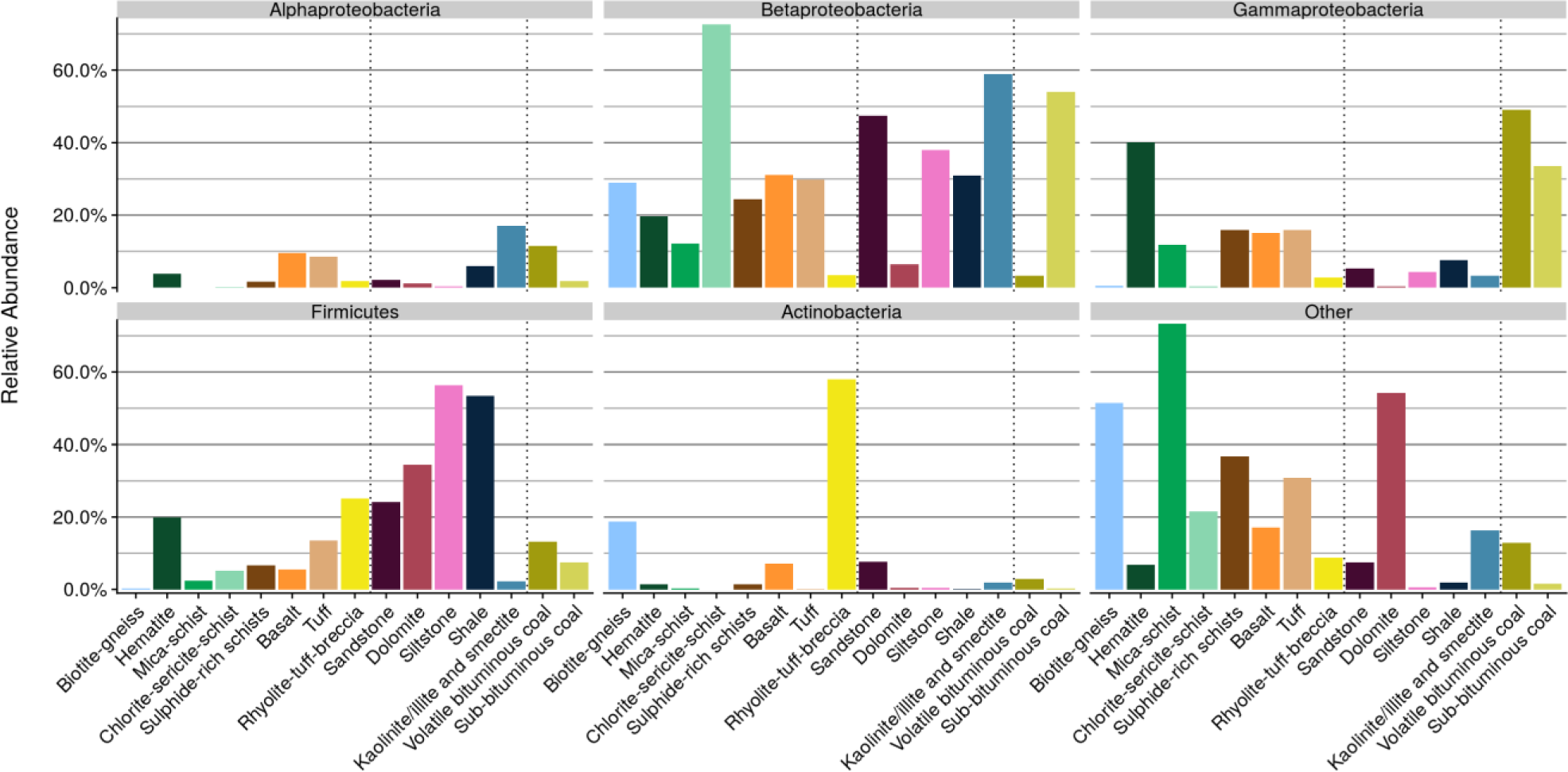
Mean relative abundances (%, y-axis) of the most abundant taxonomic groups across the dataset across all analysed aquifer lithologies (x-axis). Vertical dashed lines divide crystalline and sedimentary rocks. Coals ranks are also separated due to their higher sample contribution to the dataset.

Mean grouped proportion values indicated that Betaproteobacteria were the most abundant proteobacterial class in most host rocks, representing 26.1% of all reads in the dataset. While Betaproteobacteria accounted for 53.96% of the community profile for sub-bituminous coals, Gammaproteobacteria dominated volatile bituminous coals (49.1% of the profile, **Figure 1**). The dominance of Betaproteobacteria and Gammaproteobacteria in coals builds on culture-based evidence of widespread degradation of coal-associated complex organic compounds by these classes^56–59^.

Firmicutes were represented in large part by class Clostridia and mostly associated with sedimentary aquifers (i.e. sandstone, dolomite, siltstone, shale – **Figure 1**). This class includes ubiquitous anaerobic hydrogen-driven sulphate reducers also known to sporulate and metabolize a wide range of organic carbon compounds that have been found to dominate extremely deep subsurface ecosystems beneath South Africa and likely globally given the pervasive high levels of H_2_ in similar geologic settings ^13, 60–62^. Clostridia from metagenomes have been detected from the terrestrial deep subsurface and inferred to have the physiological capabilities needed to thrive in these environments ^20^. Adaptation to extreme environments in Clostridia is posed to be driven by varied metabolic potential, sporulation ability, and capacity for CO_2_- or sulphur-based autotrophic H_2_-dependent growth ^22, 63^.

### Lithological controls on community structure

Microbial community structure and composition in soils depend on fine-tuned geochemical, physical and hydrogeological conditions that influence microbial presence and metabolism ^64^. This relationship also appears to be reflected in the global subsurface, where host-rock lithology is evident as a primary control on community structure (**Figures 2** and **3**). Indeed, most host-rocks (10 out of 15 in this dataset) have, on average, more unique OTUs than they share with other host-rocks (**Figure 2**). Particularly, in sulphide-rich schists, 73% of the OTUs are, on average, unique to the host-rock. The role of host-rock lithology is further evidenced (**Figure 3**) as some of the host-rocks clustered at a 95% confidence interval, suggesting closely related microbial communities within similar lithologies, despite other environmental factors such as depth or location. Further, 50.6% of Jensen-Shannon distances ordinated (**Figure 3**) were significantly explained by aquifer lithology (ADONIS/PERMANOVA, F-statistic=4.65, p-value <0.001, adjusted Bonferroni correction p-value <0.001); thus showing that rock type was the primary variable defining microbial community structure. Other environmental features such as absolute sample depth and medium-scale location (i.e. state, region of the sampling site) explained only 3.08% and 2.78% of the significant metadata-driven variance in microbial community structure, respectively (ADONIS/PERMANOVA, F-statistic=3.95, 3.57 p-value <0.001, adjusted Bonferroni correction p-value <0.001). This suggests that depth-related changes in temperature and pressure are not significant controls on community structure. The relationship of community structure to hydrogeochemical parameters remains an area for future investigation – since hydrogeology and fluid geochemistry are also strongly controlled by lithology via water-rock reactions.

**Figure 2.**
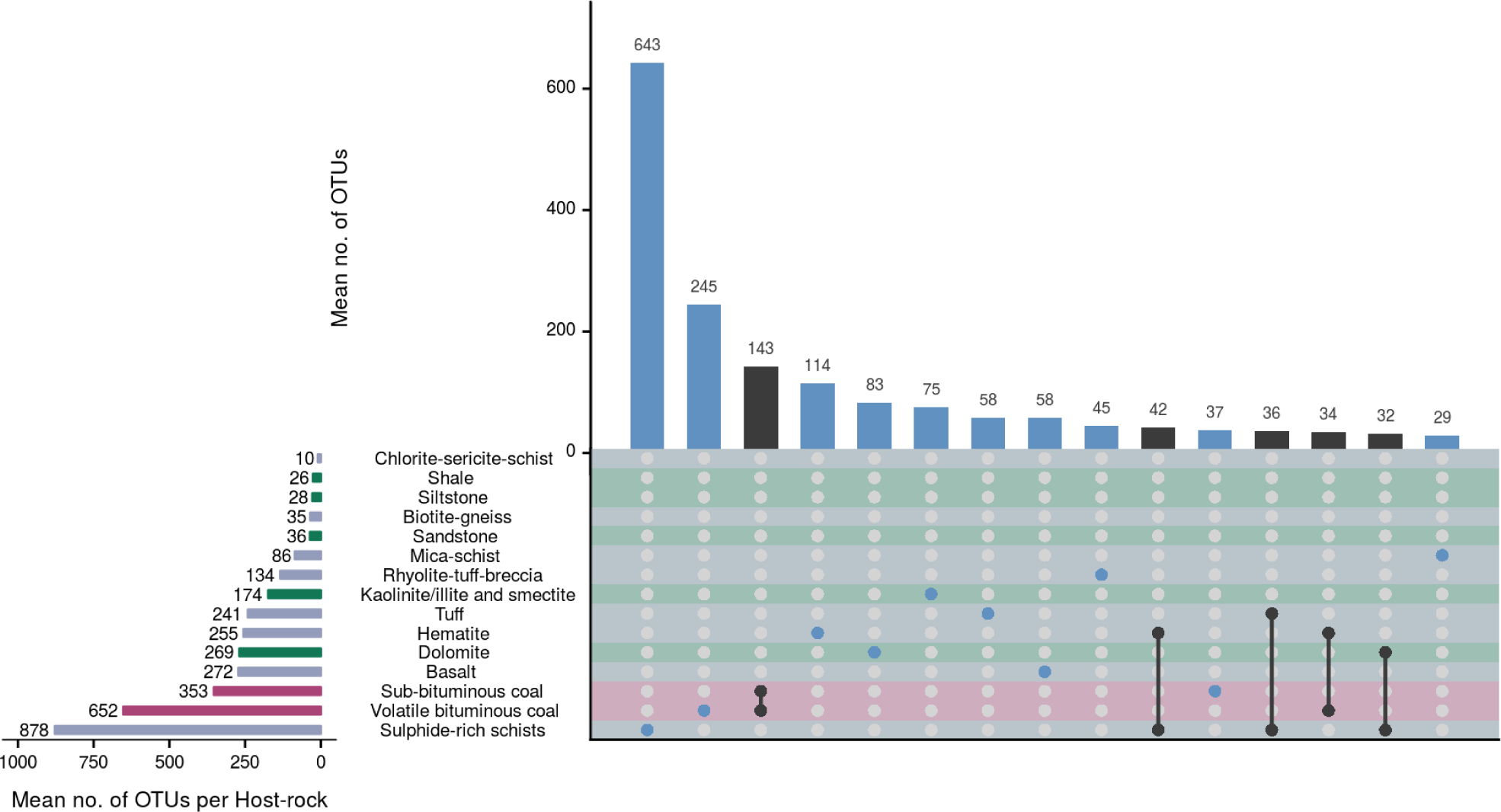
*UpsetR* plot of mean numbers of OTU interactions among rock types. Only interactions involving 25 or more OTUs on average are shown. Coloured matrix rows correspond to host-rocks, and are coloured according to rock type: blue for crystalline, green for sedimentary rocks and pink for coals, which were highlighted due to their higher sample contribution to the dataset. Columns depict OTU interactions: blue dots mark independent (mean number of non-shared OTUs) interactions and black dots connected by black lines mark shared OTUs between two or more host-rocks. Shared interactions are composed of only the host-rocks marked by dots. Vertical bars on top of the coloured matrix correspond to mean OTU numbers present in the described interactions and are coloured black or blue if depicting shared or non-shared interactions, respectively. Horizontal bars by the left of the coloured matrix depict mean total numbers of OTUs per host-rock.

**Figure 3.**
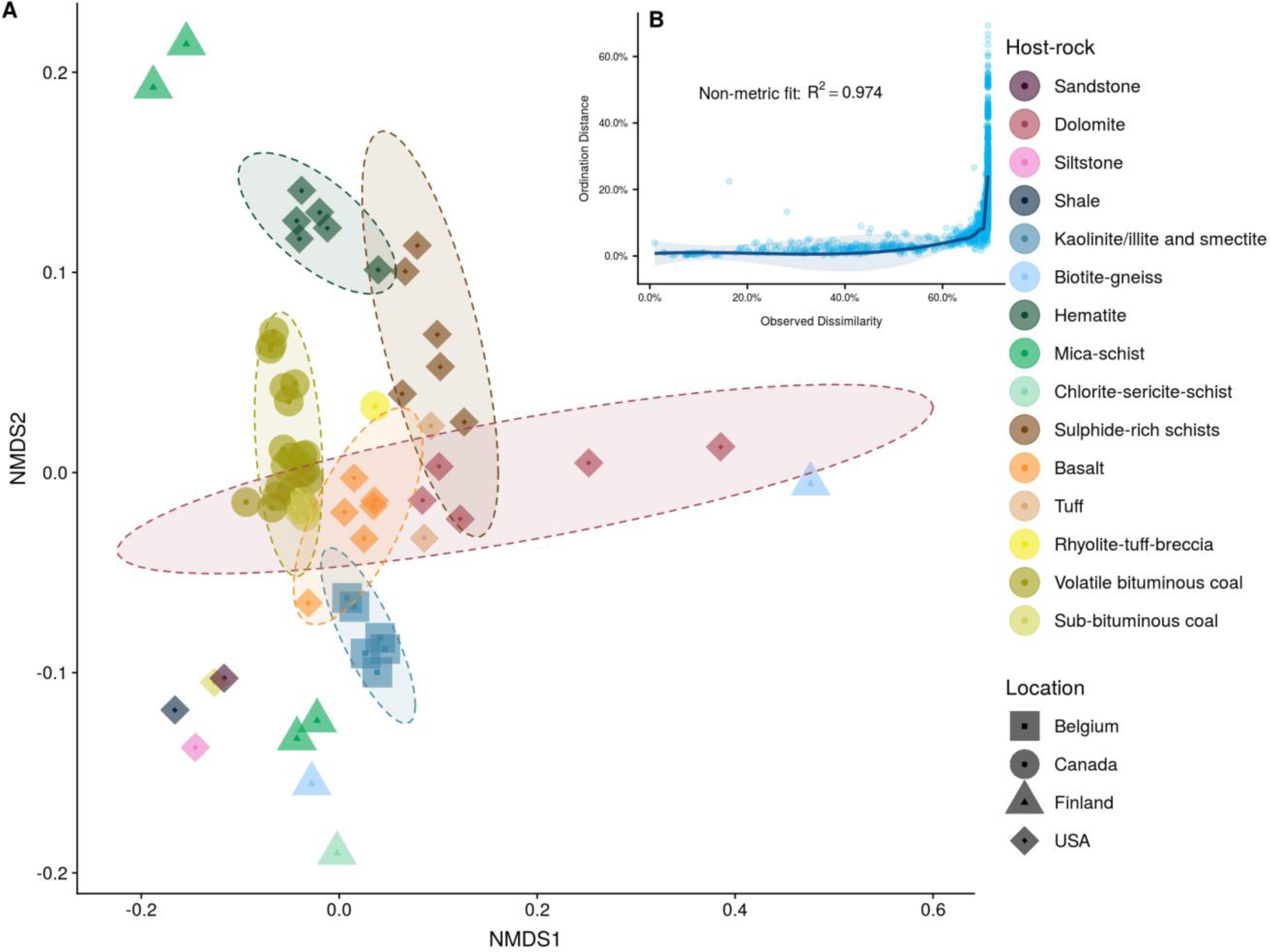
Non-metric Multidimensional Scaling (nMDS) of Jensen-Shannon distances between samples (A). Shapes correspond to different locations, whereas colours depict host-rocks targeted in this study. B depicts a Shepard’s stress plot of observed (original) dissimilarities and ordination distances. An R^2^ measure of stress is further shown for the non-metric fit of the variables. Confidence interval ellipses were plotted at a level of 95% according to host-rock.

Metadata variables that were unavailable for all samples in the dataset were excluded from the statistical analyses, thus further insights into the significance of other environmental variables was not possible. Nevertheless, this is the first large-scale evidence that deep subsurface microbial community taxonomy appears host-rock-specific. Given the importance of chemolithotrophic metabolisms in dark, subsurface environments, the unique chemical and mineralogical compositions within different aquifer lithologies impart strong controls over microbiomes associated with mineral surfaces and porewaters ^43, 62, 65–67^. Indeed, direct utilisation of mineral surfaces or dissolved species from minerals for respiration and/or metabolism has been shown to be critical in localised subsurface environments ^30, 35, 68, 69^. Due to low numbers of samples for some host-rock lithologies in this dataset (e.g. one sample each for siltstone, sandstone, shale and chlorite-sericite-schist), it is not possible to ascertain that microbiome specificity is generalisable to all deep subsurface aquifer types on Earth (see **Figure 3**). Nevertheless, this study provides the first large-scale evidence that, at a global scale, lithology surpasses depth in shaping deep subterranean microbial communities.

### A core terrestrial deep subsurface microbial community?

Analysis of prevalence across the dataset revealed that seven OTUs, all affiliated to genus *Pseudomonas*, were present in more than 25 and up to 41 samples (see **Supplementary Figure 1**, **Supplementary Table 2**). Network analysis (**Table 2**) highlighted a *Pseudomonas* OTU highly connected to other OTUs in the dataset. Further, BLAST^70^ results indicated that recovered sequences for OTUs affiliated to this genus were generally associated to marine and terrestrial soil and sediments (*cf* **Supplementary Table 4**, **Supplementary Figure 4**). Four OTUs affiliated to Burkholderiales (Betaproteobacteria), the second most prevalent order in the dataset, were also found to be connected to up to 34 other OTUs. Genus *Thauera* (Betaproteobacteria, Rhodocyclales), represented by a single OTU, was the second most central to the dataset. Finally, network and prevalence analysis highlighted the putative importance classes Betaproteobacteria and Gammaproteobacteria may have in the deep subsurface, since taxa affiliated to these were highly connected across the dataset (**Table 2**). These observations suggest genus-level taxonomy may be relevant to shaping subterranean microbial communities irrespective of host-rock lithology.

**Table 2.** Top 10 most central OTUs in the Jaccard distances network (as defined by eigenvector centrality scores, or the scored value of the centrality of each connected neighbour of an OTU) and correspondent closeness centrality (scores of shortest paths to and from an OTU to all the remainder in a network) and degree (number of directly connected edges, or OTUs) values.

| OTU ID | OTU Classification | Centrality | Closeness | Degree |
| --- | --- | --- | --- | --- |
| EF554871.1.1486 | Proteobacteria; **Gammaproteobacteria**; Pseudomonadales; Pseudomonadaceae; Pseudomonas | 1.0000000 | 2.13e-05 | 38 |
| HH792638.1.1492 | Proteobacteria; **Betaproteobacteria**; Rhodocyclales; Rhodocyclaceae; Thauera | 0.9753542 | 2.13e-05 | 36 |
| HQ681977.1.1496 | Proteobacteria; **Betaproteobacteria**; Burkholderiales; Comamonadaceae; Diaphorobacter | 0.9445053 | 2.13e-05 | 34 |
| KF465077.1.1336 | Proteobacteria; **Betaproteobacteria**; Burkholderiales; Comamonadaceae; Acidovorax | 0.8887751 | 2.13e-05 | 30 |
| JQ072853.1.1348 | Proteobacteria; **Betaproteobacteria**; Rhodocyclales; Rhodocyclaceae; Thauera | 0.8808435 | 2.13e-05 | 30 |
| KM200734.1.1449 | Proteobacteria; **Alphaproteobacteria**; Rhizobiales; Rhizobiaceae; Rhizobium | 0.8716886 | 2.13e-05 | 31 |
| KC758926.1.1392 | Proteobacteria; **Betaproteobacteria**; Burkholderiales; Comamonadaceae; Acidovorax | 0.8662805 | 2.13e-05 | 29 |
| FJ032194.1.1456 | Proteobacteria; **Betaproteobacteria**; Burkholderiales; Comamonadaceae; Rhodoferax | 0.8662805 | 2.13e-05 | 29 |
| EU771645.1.1366 | **Firmicutes**; Bacilli; Bacillales; Planococcaceae; Planomicrobium | 0.8476970 | 2.13e-05 | 30 |
| JN245782.1.1433 | Proteobacteria; **Alphaproteobacteria**.; Rhodobacterales; Rhodobacteraceae; Defluviimonas | 0.8356655 | 2.13e-05 | 29 |

The metabolic plasticity of Pseudomonadales and Burkholderiales orders has been demonstrated ^71–74^ and may be a catalyst for their apparent centrality across the terrestrial deep subsurface microbiomes analysed in this study. These bacterial orders may represent important keystone taxa in microbial consortia responsible for providing key substrates to other colonizers in deep subsurface environments ^75, 76^. In particular, given the number of highly central *Pseudomonas*-affiliated OTUs and the prevalence of this genus in the dataset, we suggest that this genus may be key in establishing conditions for microbial colonization in many terrestrial subsurface environments. Genus *Pseudomonas* and possibly several members of Burkholderiales may therefore comprise an important component of the global core terrestrial deep subsurface microbial community.

### Challenges from contamination

16S rRNA gene PCR-based approaches for characterizing microbial diversity in low biomass environments benefit from the sensitivity afforded by PCR, at the cost of vulnerability to contamination ^77^. Here, we used the prominence of sequences associated with phototrophic taxa as an indicator of either ingress of surface microbiota or contamination during sample processing. The discovery of potentially photosynthetic taxa in the initial dataset, namely 46 OTUs classified as Chloroplast (Cyanobacteria) was read as a sign that further bioinformatics-driven precautions should be taken, despite recent evidence of some cyanobacterial presence in some locations within the deep subsurface ^78, 79^. Specifically, the presence of other phototrophic members of phyla Chloroflexi and Chlorobi as well as classes Rhodospirillales (Alphaproteobacteria) and Chromatiales (Gammaproteobacteria) informed the decision to filter the dataset to hold only OTUs represented by more than 500 sequences and present in at least 10 samples. Recent recommendations for quality control of 16S rRNA gene datasets also support filtering-based approaches when applied to low biomass subsurface environments ^38^. This constraint reduced the dataset to the 70 samples and 2,207 OTUs (513,929 sequences) used for the meta-analysis (**Table 1**), and also reduced the number of prospective contaminants by half, although only ∼26% of the reads associated to Chloroplast-like sequences were removed (17 OTUs, 1958 reads).

Collecting contamination-free samples from the deep subsurface is difficult but important for cataloguing the authentic microbial diversity of the terrestrial subsurface. This study follows recent recommendations for downstream processing of contaminant-prone samples originated in the deep subsurface (Census of Deep Life project - http://codl.coas.oregonstate.edu/), where physical, chemical and biological, but also *in-silico* bioinformatics strategies to prevent erroneous conclusions have been highlighted ^38, 80, 81^. This study also follows frequency-based OTU filtration techniques similar to those recommended in Sheik *et al.* (2018) ^38^ and designed to remove possible contaminants introduced during sampling or during the various steps related to sample processing. The pre-emptive quality control steps hereby undertaken support a non-contaminant origin for taxa analysed in this dataset following careful in-field and laboratorial contamination-aware procedures carried out in each study. As such, the predominance of typically contaminant taxa affiliated *e.g.* to genus *Pseudomonas* was accepted as a true trend in the microbial ecology of the terrestrial deep subsurface.

No evidence was found for DNA extraction and PCR procedures significantly affecting microbial community structure in this meta-analysis (6.01% of the microbial community structure cumulatively [ADONIS/PERMANOVA, F-statistic=3.85, 3.23, p-value <0.01, adjusted Bonferroni correction p-value <0.001] vs. 50.6% from host rock lithology). In spite of this, a general convergence in DNA extraction methods would help further reduce methodology based variation and to standardize downstream analysis of deep subsurface microbial datasets ^82^, despite the practical challenges of each host-rock matrix and local geochemical conditions.

In the near future, the advent of recently developed techniques for primer bias-free long read 16S rRNA and 16S rRNA-ITS gene amplicon long-read-based sequencing may initiate a convergence of molecular methods from which the deep subsurface microbiology community would benefit greatly ^83, 84^. The future of large-scale, collaborative deep subsurface microbial diversity studies should encompass not only an effort towards standardization of several molecular biology techniques but also the long-term archival of samples ^85^. This will permit re-analyses using updated or unified methods after collection, where methodological variations would be controlled, and robust conclusions would more easily be achieved.

## Conclusions

A global scale meta-analysis addressing the available 16S rRNA gene-based studies of the deep terrestrial subsurface revealed the dominance of Betaproteobacteria, Gammaproteobacteria and Firmicutes across this biome. Further, aquifer lithology was identified as the main driver of deep subterranean microbial communities. Depth and location were not significant controls of microbial community structure at this scale. Finally, evidence for a core terrestrial deep subsurface microbiome population was recognised through the prevalence and centrality of genus *Pseudomonas* (Gammaproteobacteria) and several other genera affiliated to class Betaproteobacteria. The adaptable metabolic capabilities associated to the above-mentioned taxa may be critical for colonizing the deep subsurface and sustaining communities. The terrestrial deep subsurface is a hard-to-reach complex ecosystem crucial to global biogeochemical cycles. This study attempts to consolidate a global-scale understanding of taxonomical trends underpinning terrestrial deep subsurface microbial ecology and geomicrobiology.

## Methodology

### Data acquisition

The Sequence Read Archive database of the National Center for Biotechnology Information (SRA-NCBI) was queried for 16S rRNA-based deep subsurface datasets (excluding marine and ice samples, as well as any human-impacted samples); available studies, were downloaded using the SRA Run Selector. Studies were selected considering the metadata and information on sequencing platform used – i.e., only samples derived from 454 pyrosequencing and Illumina sequencing were considered. Analysis of related literature resulted in the detection of other deposited studies previous search efforts in NCBI-SRA failed to detect. Further private contacts allowed access to unpublished data included in this study. The final list of NCBI accession numbers, totalling 222 samples, was downloaded using *fastq-dump* from the SRA toolkit (https://www.ncbi.nlm.nih.gov/sra/docs/toolkitsoft/).

As seen in **Table 1**, required metadata included host-rock lithology, general and specific geographical locations, depth of sampling, DNA extraction method, sequenced 16S rRNA gene region and sequencing method. Any samples for which the above-mentioned metadata could not be found were discarded and not considered for downstream analyses.

### Pre-processing of 16S rRNA gene datasets

A customised pipeline was created in *bash* language making use of *python* scripts developed for QIIME v1.9.1 ^86^, to facilitate bioinformatic analyses in this study (see https://github.com/GeoMicroSoares/mads_scripts for scripts). Briefly, demultiplexed FASTQ files were processed to create an OTU table. Quality control steps involved trimming, quality-filtering and chimera checking by means of USEARCH 6.1 ^87^. Sequence data that passed quality control were then subjected to closed-reference (CR) OTU-picking on a per-study basis using UCLUST ^87^ and reverse strand matching against the SILVA v123 taxonomic references (https://www.arb-silva.de/documentation/release-123/). Closed-reference OTU picking excludes OTUs whose taxonomy has not been found in the 16S rRNA gene database used. Although this limits the recovery of prokaryotic diversity to the recorded in the database, cross-study comparisons of microbial communities generated by different 16S rRNA gene primers are made possible. This conservative approach classified OTUs in each study individually to the common 16S rRNA gene reference database the merge of all classification outputs. A single BIOM (Biological Observation Matrix) file was generated using QIIME’s *merge_otu_tables.py* script. The BIOM file was then filtered to exclude samples represented by less than 2 OTUs using *filter_samples_from_otu_table.py,* as well as OTUs represented by one sequence (singleton OTUs) by using *filter_otus_from_otu_table.py*. In an attempt to reduce the impacts of potential contaminant OTUs from the dataset, the post-singleton filtered dataset was further filtered to include only OTUs represented by at least 500 sequences and present in at least 10 samples overall using *filter_otus_from_otu_table.py*.

### Data analysis

All downstream analyses were conducted using the *phyloseq* (https://github.com/joey711/phyloseq) package within R, which allowed for simple handling of metadata and taxonomy and abundance data ^88–90^. Merged and filtered BIOM files were imported into R using internal *phyloseq* functions, which allowed further filtering, transformation and plotting of the dataset (see https://github.com/GeoMicroSoares/mads_scripts for scripts).

Briefly, following a general assessment of the number of reads across samples and OTUs, *tax_glom* (*phyloseq*) allowed the agglomeration of the OTU table at phylum-level. For the metadata category-directed analyses, function *merge_samples* (*phyloseq*) created averaged OTU tables, which permitted testing of hypotheses for whether geology or depth had significant impacts on microbial community structure and composition. Computation of a Jensen-Shannon divergence PCoA (Principal Coordinate Analysis) was achieved with ordinate (*phyloseq*) which makes use of metaMDS (*vegan*) ^91, 92^. All figures were plotted making use of the *ggplot2* R package (https://github.com/tidyverse/ggplot2), except for the UpsetR plot in **Figure 2**, which was plotted with package *UpsetR* (https://github.com/hms-dbmi/UpSetR).

## Acknowledgements

The work was funded by a National Research Network for Low Carbon Energy and Environment (NRN-LCEE) grant to ACM and AE from the Welsh Government and the Higher Education Funding Council for Wales (Geo-Carb-Cymru). Deep borehole samples from Nevada and California, USA (e.g. Nevares Deep Well 2 and BLM-1) were obtained with help in the field from Alexandra Wheatley, Jim Bruckner, Jenny fisher and Scott Hamilton-Brehm, and technical assistance and funding from the US Department of Energy’s Subsurface Biogeochemical Research Program, the Hydrodynamic Group, LLC, the Nye County Nuclear Waste Repository Program Office (NWRPO), the US National Park Service, and Inyo Country, CA. Samples from a mine in Northern Ontario Canada were obtained with funding from the Natural Sciences and Engineering Research Council of Canada and the assistance of Thomas Eckert, and Greg Slater of McMaster University. The Census of Deep Life (CoDL) and Deep Carbon Observatory (DCO) projects are acknowledged for a range of studies used in this analysis, as well as the sequencing team at the Marine Biological Laboratory (MBL). Disclaimer: Any use of trade, firm, or product names is for descriptive purposes only and does not imply endorsement by the U.S. Government.

## Author Contributions

ARS developed the methodology, collated and analysed the data, and wrote the manuscript. AE and AM conceived the study, supervised AS and helped write the manuscript. Other authors provided data from field sites used in the global meta-analysis. All authors contributed, edited and approved the final manuscript.

## Supplementary Text

### Depth weakly controls microbial community structure

Life at extreme depths is yet to be analysed at a deep genomic level, but microbial cells have been discovered at depths down to 3,6 km in the terrestrial crust ^1, 2^. Although cell numbers tend to decrease with depth in both crystalline and sedimentary rock in the continental crust, not much is known regarding large-scale taxonomic trends ^2^. No significant correlations were found for the presence of the most abundant clades in the dataset and depth, being Actinobacteria the only major taxonomic group to have a positive, albeit weak, correlation to depth (Pearson’s *r* = 0.42, p<0.01, **Figure S2**). Actinobacteria have already been detected at great depths in both the continental and oceanic crusts and some of its members have further been reported to hold ancestral genes for pyruvate oxidoreductase activity, which could potentially propel microbial growth at higher temperatures ^3, 4^. Proportions of Beta- and Gammaproteobacteria decreased with depth (Pearson’s *r* = −0.29, Pearson’s *r* = −0.093), and no other major clades were shown to correlate. More data needs to be generated to better investigate which, if any, taxonomic groups prefer deeper terrestrial environments. Biochemical limitations to life such as racemization rates of organic acids with depth and temperature will surely select for adaptable taxa and possibly create a mostly depth-defined gradient of representation of adapted extremophilic clades ^5^.

### 99% closed-reference OTU analysis

A 99% similarity closed-reference OTU-picking strategy to further minimize potential contamination showed a ∼10-fold decrease in the number of OTUs and read numbers (1,065 OTUs and 70,527 reads were left). Further, this reduced the number of retrieved samples to 93.

Following the previously described filtering steps (hold only OTUs represented by more than 500 sequences and present in at least 10 samples), 335 OTUs (67,151 reads) associated to 14 samples were left. For this reduced dataset, 2 OTUs associated to Chloroplast were found, represented by 115 sequences in total. The very reduced number of samples, OTUs an total reads left in the datasset caused the 99% OTU approach to be discontinued.

## Supplementary Methodology

### Phylogeny of Pseudomonas representative sequences

Representative 16S rRNA gene sequences for *Pseudomonas* OTUs in the dataset were isolated by retrieving OTU IDs affiliated to this genus in the final dataset using the *subset_taxa* function within *phyloseq*^6^. The SILVA 123 database was then queried against the list of OTU IDs and the results deposited in a FASTA file. Outgroup sequences of genus *Sphingomonas* (Proteobacteria, Alphaproteobacteria, Sphingomonadales, Sphingomonadaceae) were obtained directly from the SILVA database (https://www.arb-silva.de/) and added along with the retrieved *Pseudomonas* 16S rRNA gene sequences to a final FASTA file.

Using MEGA7^7^, an alignment was performed using the MUSCLE^8^ algorithm and 8 iterations of the UPGMB (combines Neighbour-Joining^9^ and UPGMA - Unweighted Pair Group Method with Arithmetic mean) clustering method. An optimal Neighbour-Joining^9^ tree (*cf.* **Supplementary Figure 4**) was then created using 500 bootstrapped replicates and the Maximum Composite Likelihood (MCL)^10^ method to calculate evolutionary distances.

---

**Supplementary Figure S1.**
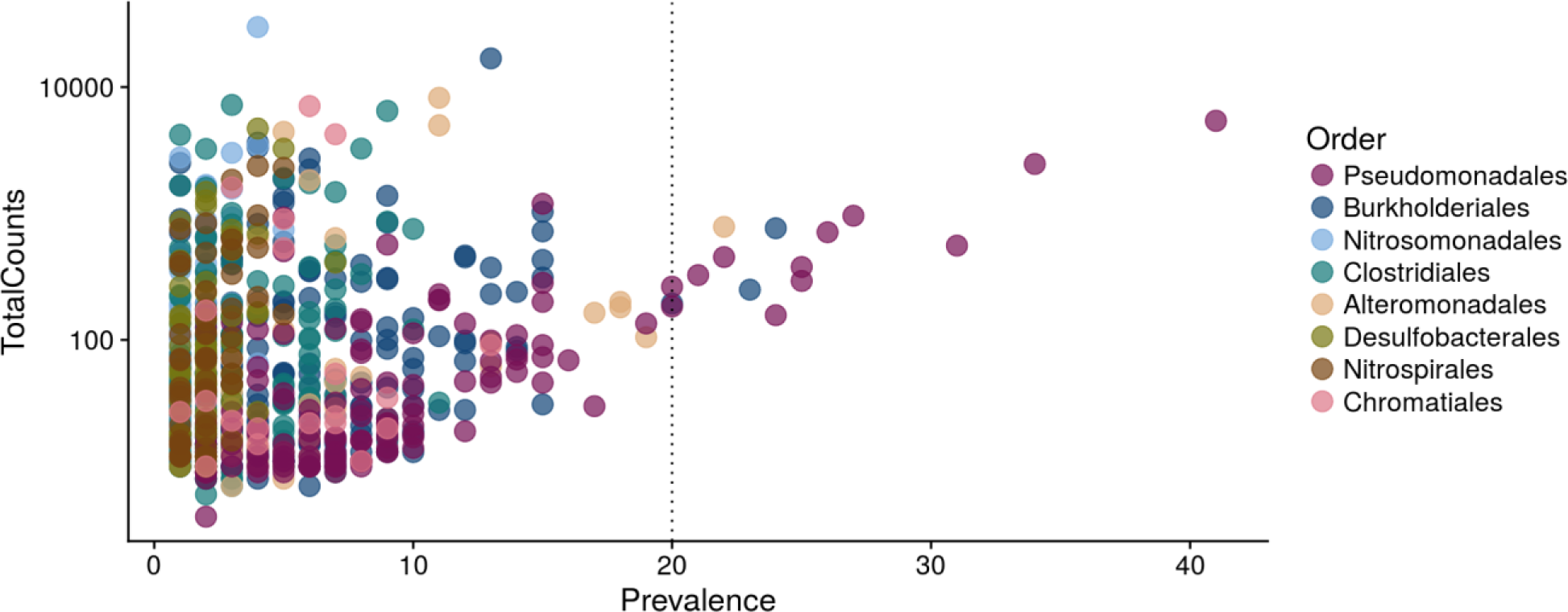
Prevalence (number of samples an OTU is present in, x-axis) of OTUs across the dataset and associated reads (y-axis). Colours depict classification of OTUs at order level. Vertical line crosses 20 samples in the x-axis to highlight OTUs present in 20 or more samples.

**Supplementary Figure S2.**
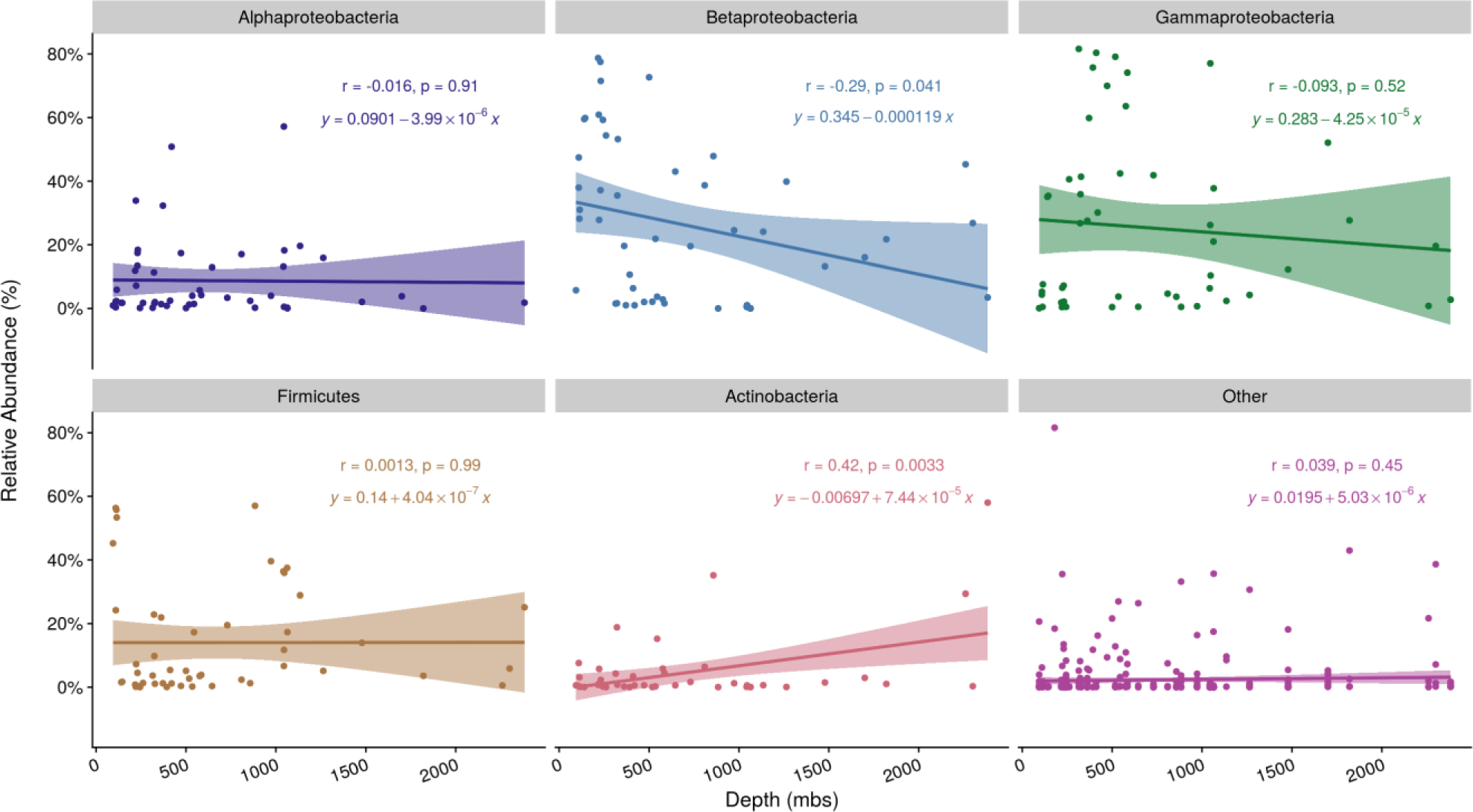
Correlations between relative abundance of OTUs (%, y-axis) associated to the most abundant taxonomic groups across the dataset and depth (meters below surface, x-axis). Regression lines follow the linear model and shading around lines corresponds to the 95% confidence interval. Annotations in plot facets indicate the associated Pearson correlation coefficient, its corresponding p-value and the fitted linear model equation. Each point represents an OTU associated to the taxonomic group in each facet at a certain depth - a same OTU may be depicted more than one time.

**Supplementary Figure S3.**
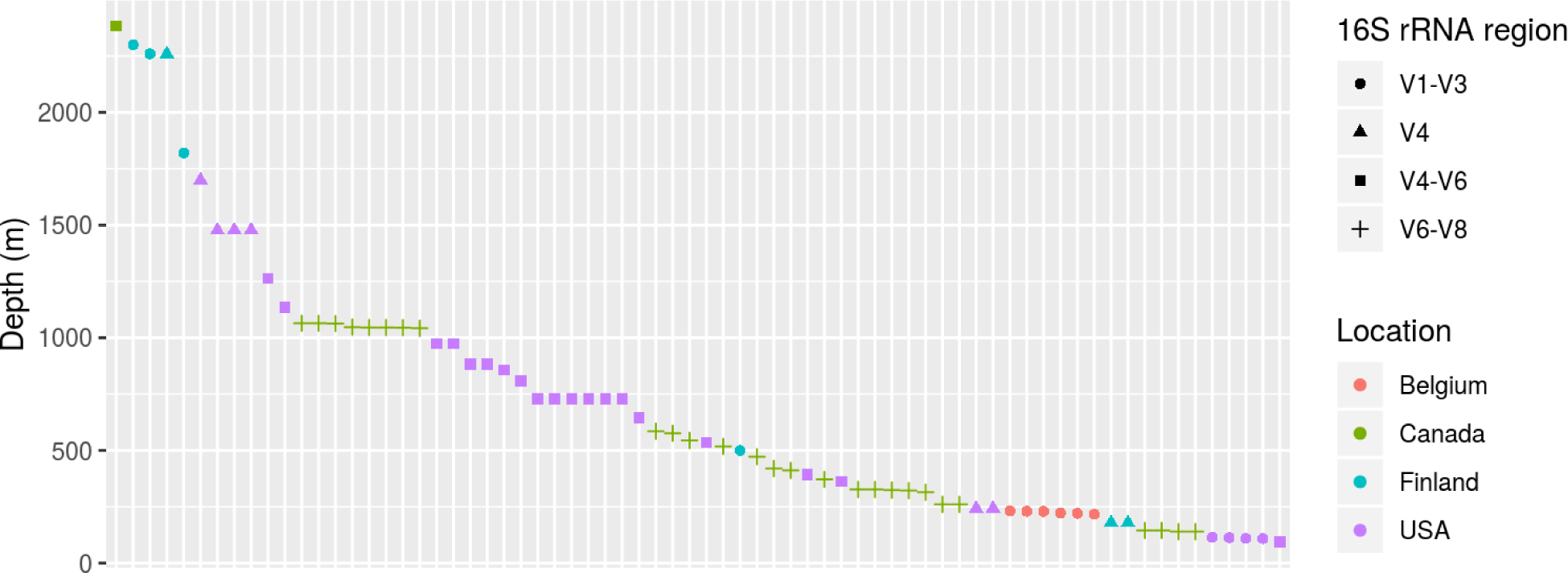
Distribution of samples in the final dataset across depth, colourised by general location. Shapes indicate 16S rRNA gene region utilised for that study.

**Supplementary Figure S4.**
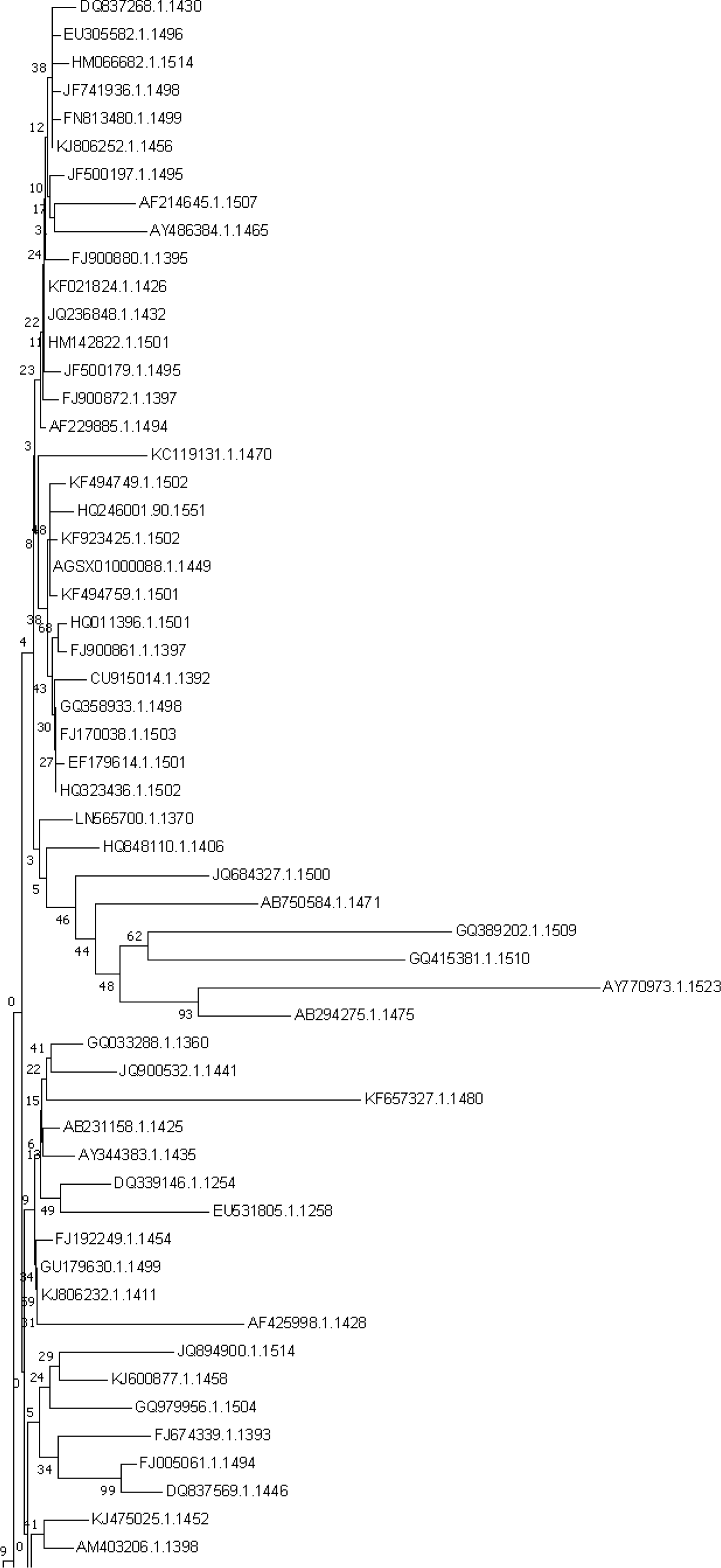
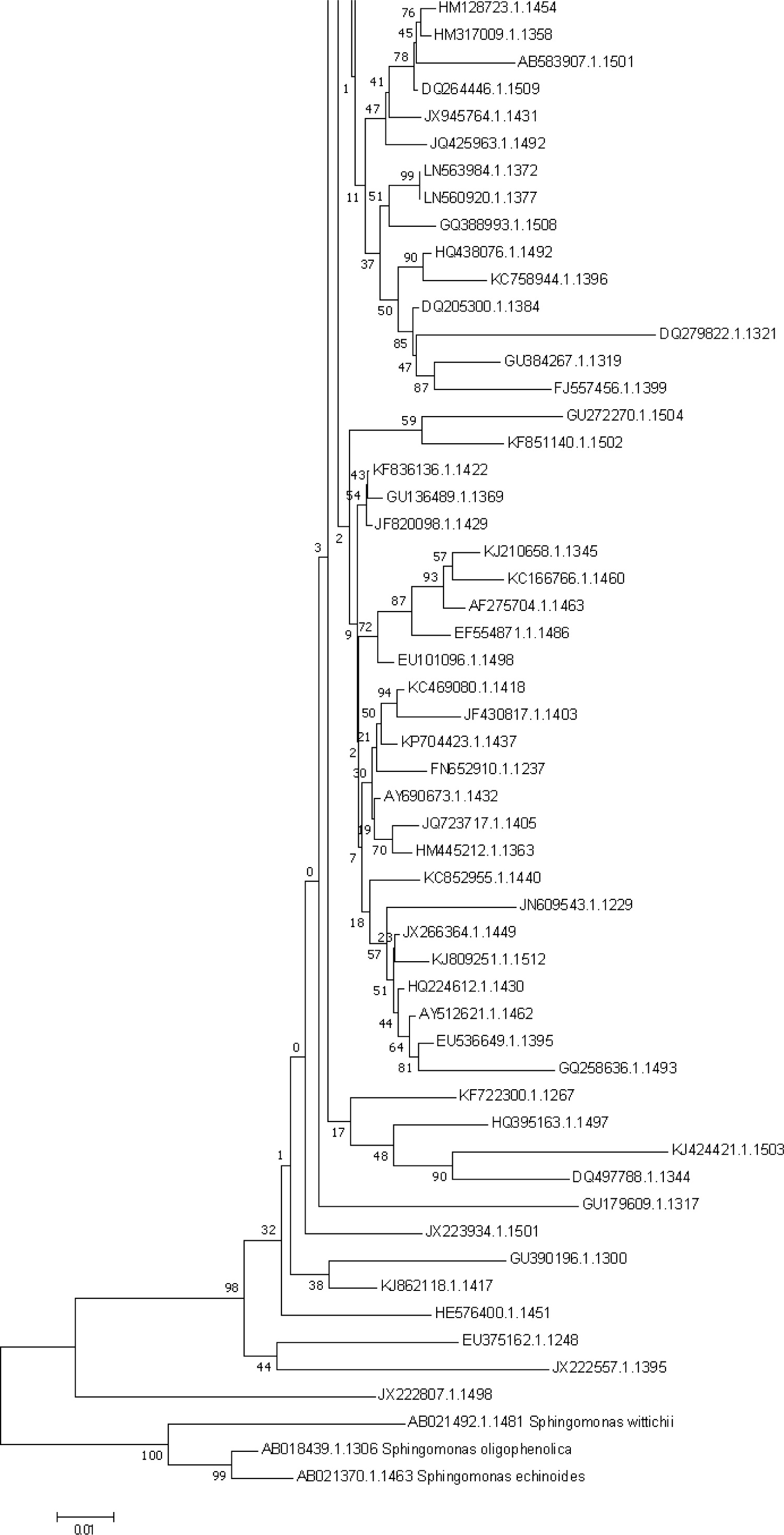
Optimal Neighbor-Joining tree of representative sequences for *Pseudomonas* OTUs in the dataset following MUSCLE aligment. OTU IDs are shown in the end of each branch and outgroup sequences of genus *Sphingomonas* identified by their taxonomic affiliation. The percentage of bootstrap test replicates are shown next to tree branches and scale for MCL evolutionary distances is shown in the bottom.

**Table S1.** Top 10 OTU network interactions ordered by edge betweenness (number of shortest paths going through an edge - OTU/OTU interactions) values as per the calculated Jaccard distances.

| OTU 1 Classification | OTU 2 Classification | Betweenness |
| --- | --- | --- |
| Gammaproteobacteria, Chromatiales, Chromatiaceae, Rheinheimera | Gammaproteobacteria, Chromatiales, Chromatiaceae, Rheinheimera | 1686 |
| Gammaproteobacteria, Chromatiales, Chromatiaceae, Rheinheimera | Gammaproteobacteria, Pseudomonadales, Moraxellaceae, Acinetobacter | 1656.17 |
| Gammaproteobacteria, Chromatiales, Chromatiaceae, Rheinheimera | Gammaproteobacteria, Alteromonadales, Alteromonadaceae, Alishewanella | 885.1 |
| Gammaproteobacteria, Pseudomonadales, Pseudomonadaceae, Pseudomonas | Gammaproteobacteria, Chromatiales, Chromatiaceae, Rheinheimera | 749 |
| Gammaproteobacteria, Pseudomonadales, Pseudomonadaceae, Pseudomonas | Gammaproteobacteria, Chromatiales, Chromatiaceae, Rheinheimera | 618 |
| Betaproteobacteria., Rhodocyclales, Rhodocyclaceae, Azoarcus | Betaproteobacteria, Rhodocyclales, Rhodocyclaceae, uncultured | 588 |
| Firmicutes, Clostridia, Clostridiales, Peptostreptococcaceae, Peptoclostridium | Firmicutes, Clostridia, Clostridiales, Peptostreptococcaceae, Intestinibacter | 588 |
| Gammaproteobacteria, Pseudomonadales, Pseudomonadaceae, Pseudomonas | Alphaproteobacteria, Rhodobacterales, Rhodobacteraceae, Paracoccus | 563.83 |
| Gammaproteobacteria, Chromatiales, Chromatiaceae, Rheinheimera | Gammaproteobacteria, Alteromonadales, Alteromonadaceae, Alishewanella | 499.26 |
| Gammaproteobacteria, Pseudomonadales, Moraxellaceae, Acinetobacter | Gammaproteobacteria, Pseudomonadales, Moraxellaceae, Acinetobacter | 495 |

**Table S2.** SILVA 123 taxonomic affiliations of OTUs present in 20 or more samples. Prevalence is defined as the number of samples an OTU is present in.

| Taxa ID | Taxa Classification | Prevalence |
| --- | --- | --- |
| JQ236848.1.1432 | Proteobacteria, Gammaproteobacteria, Pseudomonadales, Pseudomonadaceae, Pseudomonas | 41 |
| HM142822.1.1501 | Proteobacteria, Gammaproteobacteria, Pseudomonadales, Pseudomonadaceae, Pseudomonas | 34 |
| JX266364.1.1449 | Proteobacteria, Gammaproteobacteria, Pseudomonadales, Pseudomonadaceae, Pseudomonas | 31 |
| KJ475025.1.1452 | Proteobacteria, Gammaproteobacteria, Pseudomonadales, Pseudomonadaceae, Pseudomonas | 27 |
| FJ192249.1.1454 | Proteobacteria, Gammaproteobacteria, Pseudomonadales, Pseudomonadaceae, Pseudomonas | 26 |
| HQ848110.1.1406 | Proteobacteria, Gammaproteobacteria, Pseudomonadales, Pseudomonadaceae, Pseudomonas | 25 |
| KC852955.1.1440 | Proteobacteria, Gammaproteobacteria, Pseudomonadales, Pseudomonadaceae, Pseudomonas | 25 |
| HM773515.1.1498 | Proteobacteria, Betaproteobacteria, Burkholderiales, Comamonadaceae, uncultured | 24 |
| AB231158.1.1425 | Proteobacteria, Gammaproteobacteria, Pseudomonadales, Pseudomonadaceae, Pseudomonas | 24 |
| JX222276.1.1475 | Proteobacteria, Betaproteobacteria, Burkholderiales, Comamonadaceae, Variovorax | 23 |
| EU841498.1.1443 | Proteobacteria; Gammaproteobacteria; Alteromonadales, Alteromonadaceae, Alishewanella | 22 |
| EU305582.1.1496 | Proteobacteria, Gammaproteobacteria, Pseudomonadales, Pseudomonadaceae, Pseudomonas | 22 |
| FJ900880.1.1395 | Proteobacteria, Gammaproteobacteria, Pseudomonadales, Pseudomonadaceae, Pseudomonas | 21 |

**Table S3.** Metadata table with complete details for all studies utilized.

| SRA Accession | Depth gradient<br>(m) | DNA Extraction method | Sample Type | 16S region | Sequencing<br>method | Project |
| --- | --- | --- | --- | --- | --- | --- |
| PRJNA262938 | 243.84-1478.28 | Mod. Phenol chloroform | Groundwater | V4 | 454 |  |
| PRJNA268940 | 94-2383 | MoBio UltraClean Microbial DNA isolation kit | Groundwater | V4-V6 | 454 | VAMPS |
| PRJNA248749 | 730 | MoBio PowerWater | Groundwater | V4-V6 | 454 | VAMPS |
| PRJNA251746 | 393-1135 | MoBio_PowerMax_Soil | Groundwater | V4-V6 | 454 | VAMPS |
| PRJNA375701 | 109-114.7 | Mod. MP-Biomedicals FastDNA Spin_Soil <sup>11</sup> | Rock + porewater | V1-V3 | 454 |  |
| NA* | 140-1064.4 | Foght *et al.* 2004 | Rock + porewater | V6-V8 | 454 | HMP |
| PRJEB1468 | 217.13-231.83 | XS buffer <sup>13</sup> | Groundwater | V1-V3 | 454 |  |
| PRJEB10822 | 180-2300 | MoBio PowerSoil | Groundwater | V4**, V1-V3 | 454 |  |

\* http://hmp.ucalgary.ca/HMP/
\* This primer pair corresponds to archaeal 16S rRNA gene primers (A344,A744, *cf.* 10.5194/bg-13-3091-2016 for further details).

**Table S4.**
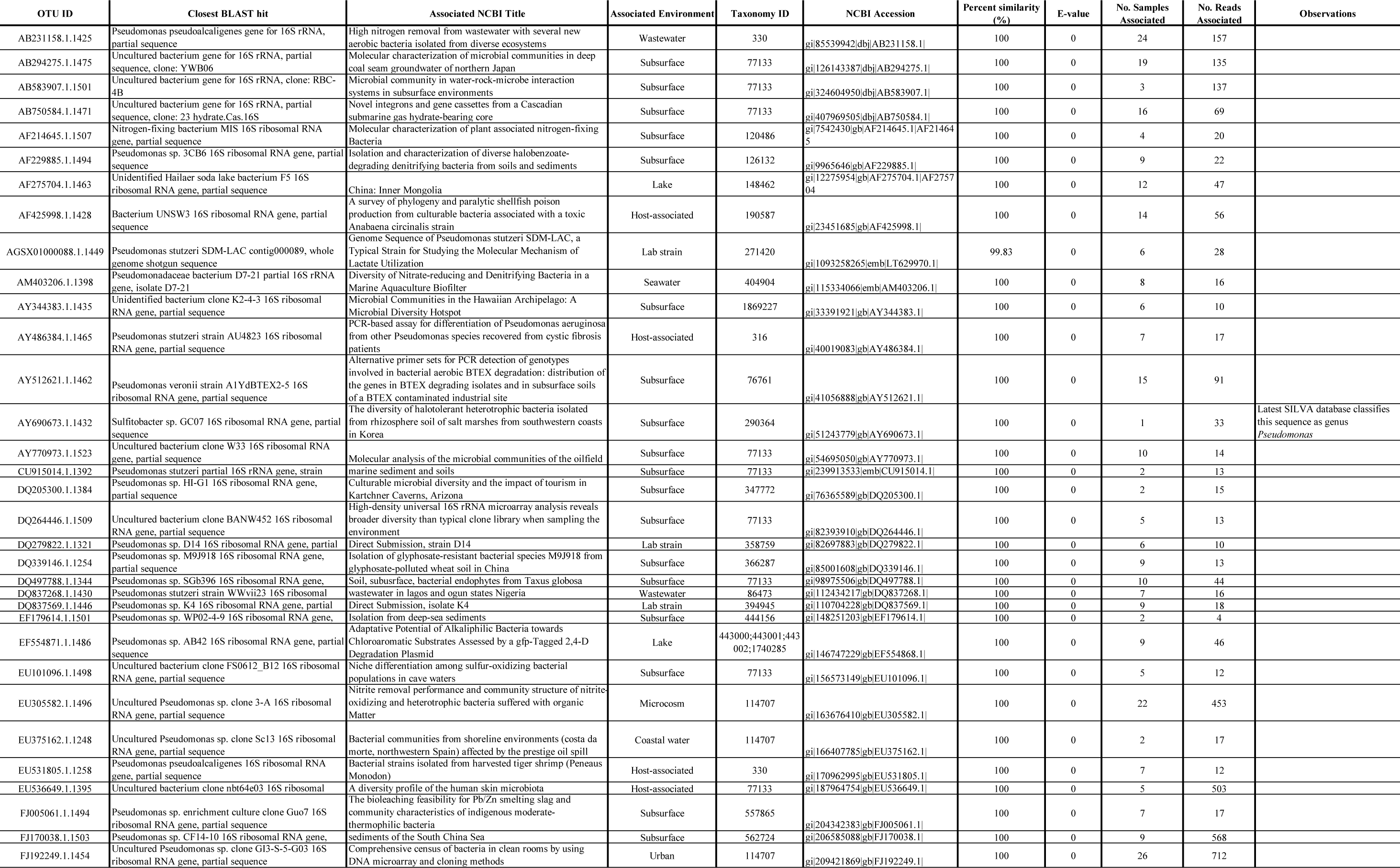
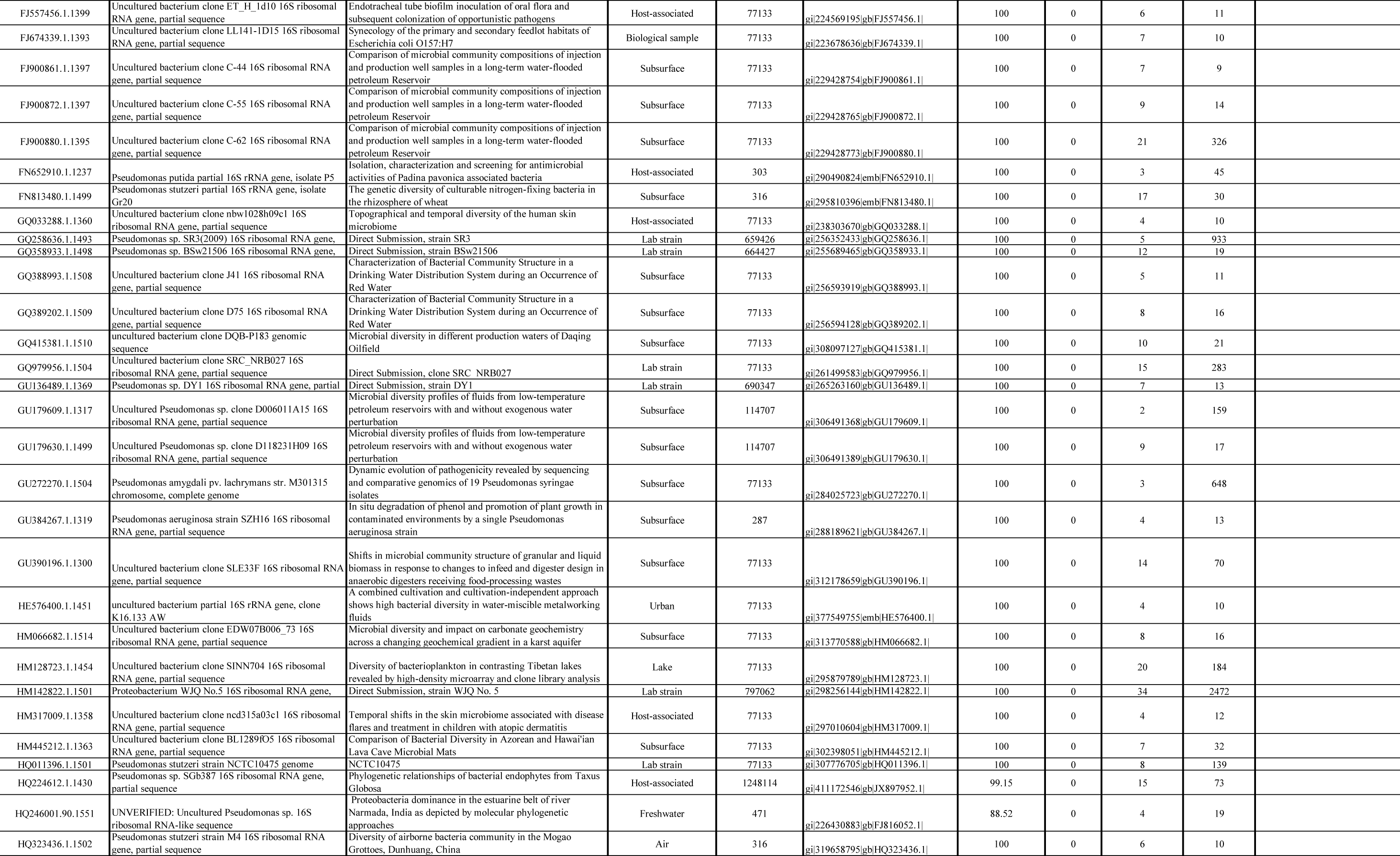
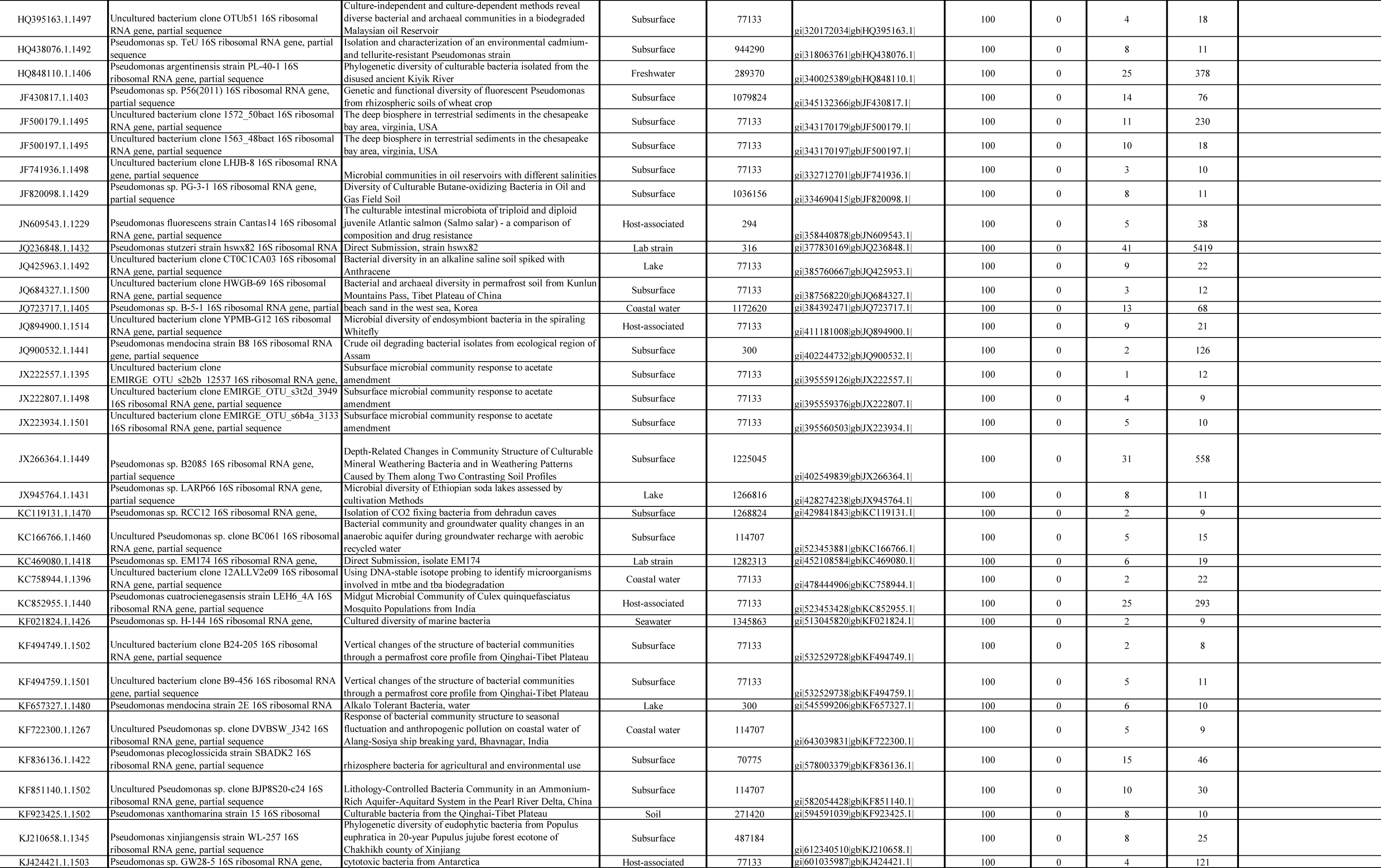
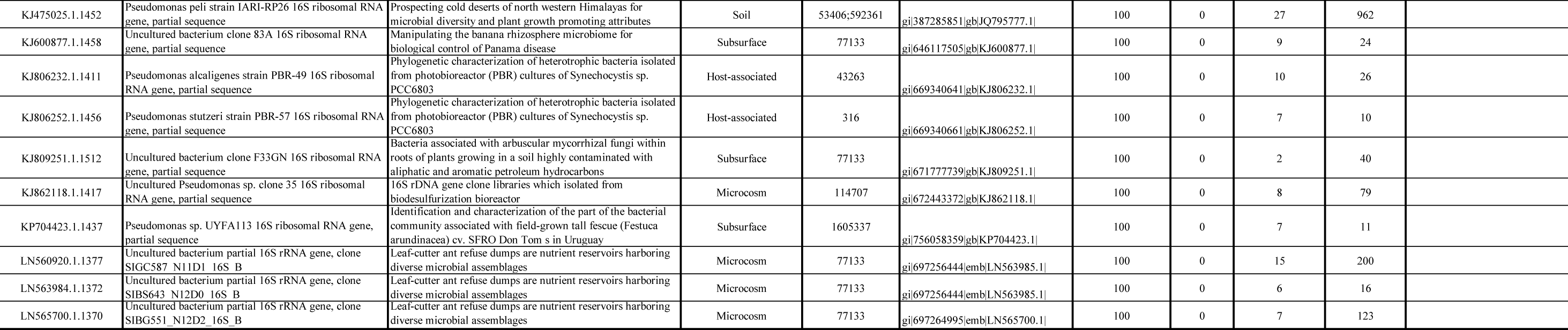
BLAST hits table for representative sequences associated to OTUs affiliated to genus Pseudomonas.

